# Less is more: uncompensated gravity torques for intuitive EMG-based assistance with a robotic exoskeleton

**DOI:** 10.1101/2025.11.21.689589

**Authors:** Lucas Quesada, Dorian Verdel, Olivier Bruneau, Bastien Berret, Michel-Ange Amorim, Nicolas Vignais

**Affiliations:** CIAMS, Université Paris-Saclay, Inria, Gif-sur-Yvette, France; LURPA, École Normale Supérieure Paris-Saclay, Gif-sur-Yvette, France; Human Robotics Group, Department of Bioengineering, Imperial College of Science, Technology and Medicine, W12 0BZ London, United-Kingdom; Univ Rennes, Inria, M2S, F-35000 Rennes, France

**Keywords:** exoskeleton, myocontroller, movement assistance, human-robot interactions, human-centered control

## Abstract

Despite extensive investigation on the use of electromyographic (EMG) activity to control active exoskeletons over the past decade, designing intuitive assistive controllers that seamlessly integrate with natural human motor control have yet to be realized. While existing EMG-based controllers often achieve substantial reduction in muscle effort, they frequently incur increased cognitive and attentional load for the user, thereby compromising the overall efficacy of the assistance. Here we introduce a novel EMG-based assistive controller founded upon neuroscience principles, specifically the observation that humans naturally exploit gravity torque to facilitate movement control. Therefore, deviating from conventional compensation strategies, our approach purposely leaves a fraction of the predicted human gravity torque uncompensated so that users can still take advantage of gravity as they would without assistance. Through a load-carrying arm movement task, we show that enabling gravity exploitation improves traditional EMG-based assistance by achieving a significant reduction in muscle effort, while concurrently yielding superior kinematic performance (i.e., faster, smoother movements) and enhanced subjective user experience. These findings demonstrate that integrating principles from neural motor control into assistive controllers allows to implement a favorable tradeoff between muscle effort reduction and functional usability.

## 1 Introduction

er limb exoskeletons are envisioned as a potential transformative technology in a large range of contexts, from the prevention of work-related musculoskeletal disorders (MSDs) [1], to improving rehabilitation protocols [2], and recovering disabled motor functions [3]. These systems are highly versatile, being able to either measure human movement with minimized impact [4–7], or flexibly provide different levels of assistance [8–11] up to fully assisted movements with minimal or null user effort [12, 13]. However, their usability in practice remains hampered by the lack of (i) non-specialized control methods, allowing to assist the user throughout multiple tasks and activities, and (ii) consideration for human motor control, both when designing the exoskeleton controller and evaluating its impact on the user, in particular when the human and exoskeleton are co-adapting through time.

In this context, assistive controllers based on the synthesis of the user’s muscle activities into assistive torques via electromyographic (EMG) signals are a promising venue. In contrast with techniques requiring to extract a human desired trajectory [11, 14–17], from a set of human demonstrations [16, 18, 19], EMG activities can be directly mapped to desired human torques implemented to control the exoskeleton. Several models, all requiring an individualized calibration, have been proposed to perform this mapping, ranging from simple linear approaches [20–23] to nonlinear musculoskeletal modeling [9,24–27], and more efficient hybrid approaches [28]. Despite requiring several process steps, inducing delays to obtain the human torque [29], EMG signals can potentially provide a predictive information regarding the user’s behavior due to the *electromechanical delay* between the muscle activation and its effect on the limb dynamics. In theory, EMG-based controllers are therefore ideal candidates to solve the limitations previously underlined.

Nonetheless, these approaches have mostly been applied to single-degree-of-freedom tasks [30–32], and controllers have often consisted in a simplistic proportional compensation of the estimated user’s torques [9, 33–35], with a validation limited to showing a decrease in measured EMG activities. Some notable alternatives have been proposed, such as (i) integrating the predicted human torques in a model predictive control scheme [30], leading to an optimal sharing of task-related efforts between the human and exoskeleton, or (ii) including an integral term in the provided assistance [32,36], leading to a performance comparable to gravity compensation methods while allowing to carry unknown masses. Importantly, the validation of these approaches remained very limited, with few or no consideration for the effects of the developed controller on human motor control and for their perception by users.

In particular, it is well known in the motor control literature that humans exploit gravity-related efforts to accelerate downwards movements and slowdown upward movements [37], allowing them to optimize their energetic expenditure [38] (see Fig. 1.a). This human adaptation to a ubiquitous component of their environment has tractable effects on their movements kinematics [39] and muscle activation patterns [40], also observable in other primates [41] and when interacting with robotic exoskeletons [10]. Given that gravity exploitation is a major feature of human volitional movement, it is currently uncertain whether maximally compensating for the user’s effort (including those related to gravity) is the best strategy to progress towards symbiotic human-exoskeleton interactions. Crucially, full effort compensation leads human users to develop a non-natural motor strategy in which they have to contract extensor muscles to initiate downward movements, whereas in natural settings they would simply relieve flexor (antigravity) muscles [42]. Therefore, we hypothesize that leaving some gravity-torques uncompensated may lead to a more intuitive and seamless assistance than when minimizing human effort during a full gravity compensation (see Fig. 1.b).

**Figure 1:**
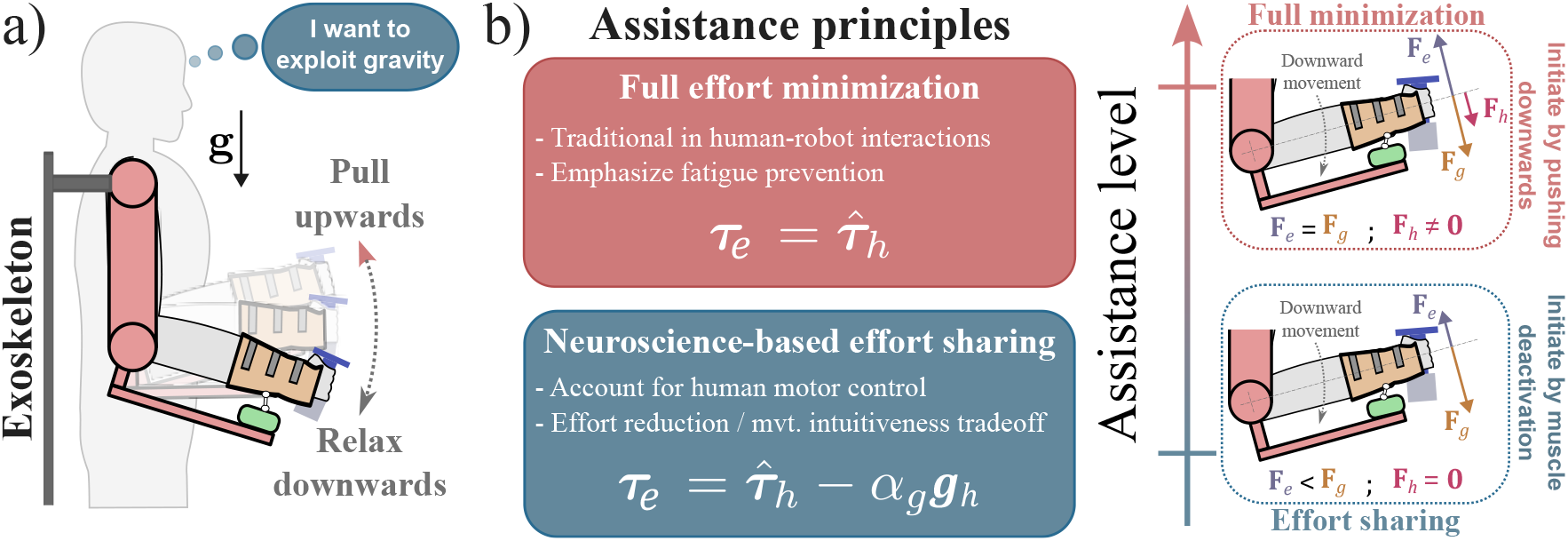
Neuroscience informed EMG-based assistance concept. **a)** When subjected to gravity, tasks requiring vertical movements warrants differentiated muscle patterns for upward and downward directions [37, 38, 40, 41]. Upward movements are initiated by a contraction of antigravity muscles (i.e. generating antigravity torques), while downward movements are initiated by a deactivation of those same muscles. **b)** Here, we compare the full effort minimization strategy–controlling the exoskeleton torques ***τ***_*e*_ to fully compensate the EMG-estimated human torques 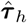 –to our neuroscience-based effort sharing strategy–leaving a fraction *α*_*g*_ of gravity-related torques uncompensated. The traditional approach aims at minimizing muscle fatigue and human energy expenditure [9], but require users to activate gravity muscles (i.e. generating torques in the same direction as gravity) to initiate downward movements. Hence, the resulting robot force **F**_*r*_ equilibrates with the gravity-related forces **F**_*g*_ in static, and the human can remain passive (i.e. **F**_*h*_ = 0 before movement). In contrast, with our method the human has to provide a torque *α*_*g*_***g***_*h*_ to maintain the static equilibrium, and downward movements are initiated by relaxing antigravity muscles.

In the present paper, we aim to provide and evaluate an EMG-based assistance strategy mitigating the abovementioned limits, i.e. (i) a non-specialized control method, usable for any task, while (ii) accounting for knowledge in human motor control, validated through an in-depth evaluation of the resulting human-machine interaction. Therefore, we will compare user efforts, movement performance, and perceptual feedback without and with an exoskeleton providing different levels of physical assistance. In particular, users interact with (i) a transparent exoskeleton, following their movements with minimized interaction efforts [7], (ii) a proportional-integral EMG-based assistance combining previous developments [9, 32], and (iii) a novel EMG-based assistance considering human motor control under gravity-related efforts (see Fig. 1.b). We aim to validate our different controllers on a multiple degrees-of-freedom task, with evolving dynamics, so as to obtain results that are as general as possible. Specifically, we ask users to reach, grab, move, and place a mass unknown to the robot on precise locations, similar to a pick-and-place task that could be encountered in logistics warehouses [43], with movements in a parasagittal plane inducing varying gravity-related efforts.

## 2 Results

We asked 10 participants to perform a mass displacement task between two locations, one above and beyond the other in a parasagittal plane (see Fig. 2b), requiring them to use elbow and shoulder flexion and extension movements. The movements to perform in the task can be described by four categories: (i) reaching towards the mass when it is placed on the further site (*Reach high*); (ii) reaching towards the start position after placing the mass on the further site (*Reach low*); (iii) moving the mass towards the closer site (*Place low*); or (iv) moving towards the further site (*Place high*). Users were asked to perform cycles or *Reach high*–*Place low* and *Place high*–*Reach low* movements without external intervention between the cycles. This implies that the controllers have to handle the transitions between movements performed with and without the mass. The exoskeleton could be controlled to provide an EMG-based assistance (EA) resulting from a proportional integral control, completely minimizing users’ efforts. Alternatively, we let a proportion of the gravity-related efforts be handled by the user by dynamically removing it from their estimated joints torques (EAG). For baseline comparisons, we also asked the participants to perform the task without the exoskeleton (NE) and with the exoskeleton controlled in transparent mode (TR).

**Figure 2:**
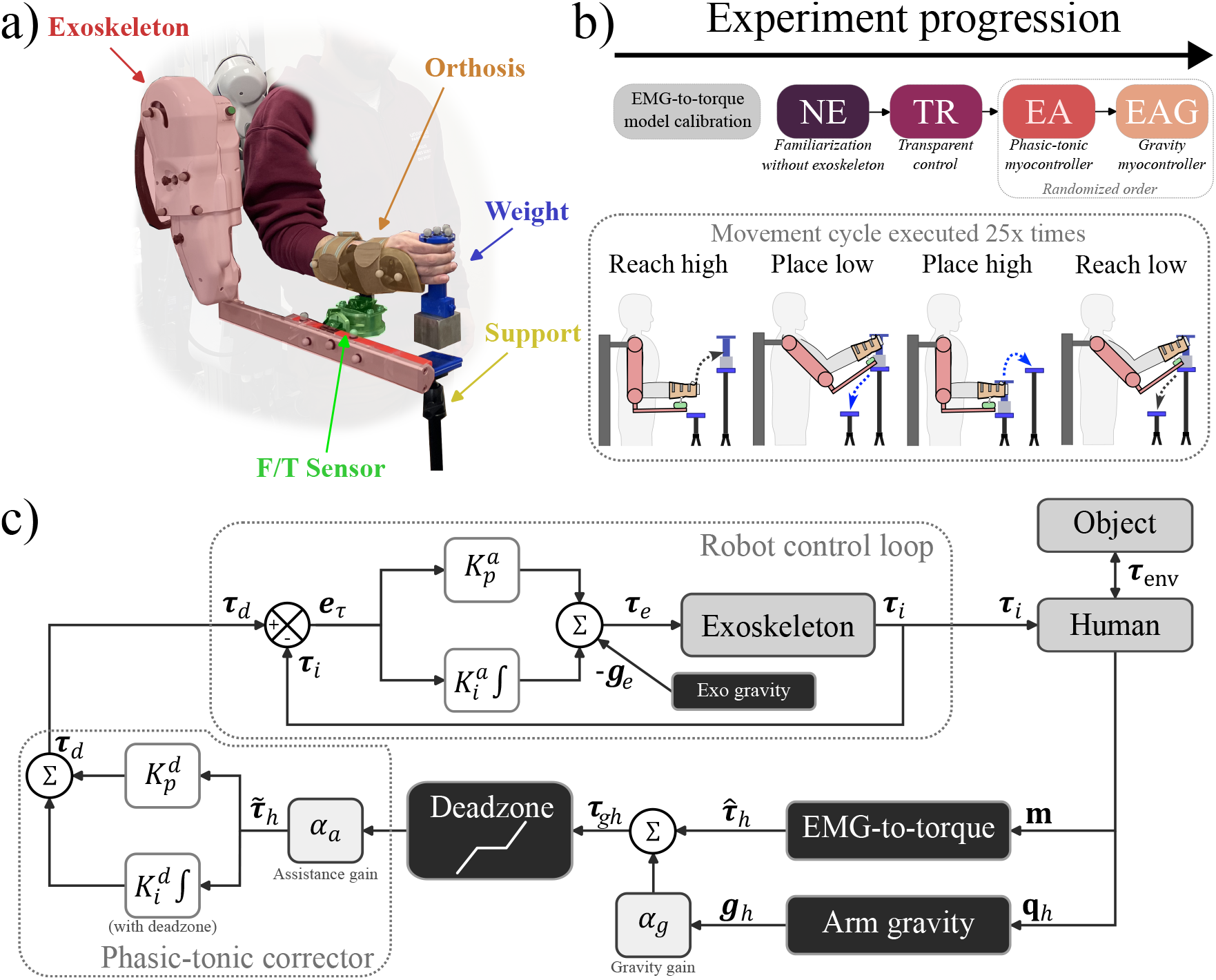
Setup, multi-DoF pick-and-place taskn, and investigated controllers. **a)** Participants were connected to the exoskeleton through an ergonomic orthosis [44,45], instrumented with a Force/Torque (FT) sensor. They carried the weight from one support to the other by grabbing it with their own hand. **b)** After calibrating our hybrid EMG-to-torque model [28], participants were asked to perform 25 cycles of mass retrieval and displacement per condition. These cycles are first executed without the exoskeleton for familiarization (NE), then with a transparent controller (TR), and finally with the myocontrollers (EA and EAG) in a randomized order. **c)** The exoskeleton was controlled to compensate its own weight ***g***_*e*_ and minimize the torque error *e*_*τ*_ –between the measured ***τ***_*i*_ and desired ***τ***_*d*_ interaction torques–via a proportional-integral admittance loop using the FT sensor with gains 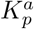 and 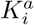 respectively. Human muscle activities **m**, resulting from the interaction with the robot ***τ***_*i*_ and environment ***τ***_env_, allowed to estimate human torque inputs 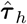. Gravity torques were estimated from the exoskeleton’s joints angles and anthropometric tables, allowing to implement the EAG controller. For all EMG-based assistance conditions, the torques ***τ***_*gh*_ were filtered with a deadzone and multiplied by an assistance gain *α*_*a*_ (set to *α*_*a*_ = 0 in TR) to obtain the human intent torque to compensate 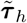. The exoskeleton desired torques were then computed using a proportional-integral scheme with gains 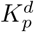 and 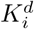, allowing to memorize recent human inputs to compensate for (a fraction of) weight when muscle activity diminished in static.

### 2.1 EMG-based assistance reduces effort

The primary aim of assistive controllers developed in the present paper is to reduce human effort. Consequently, the performance of EA and EAG was first assessed in terms of human energetic expenditure estimated from (i) the absolute mechanical work in the different phases of the task, and (ii) the level of EMG activity. The summary of this first analysis is described in Fig. 3a.

**Figure 3:**
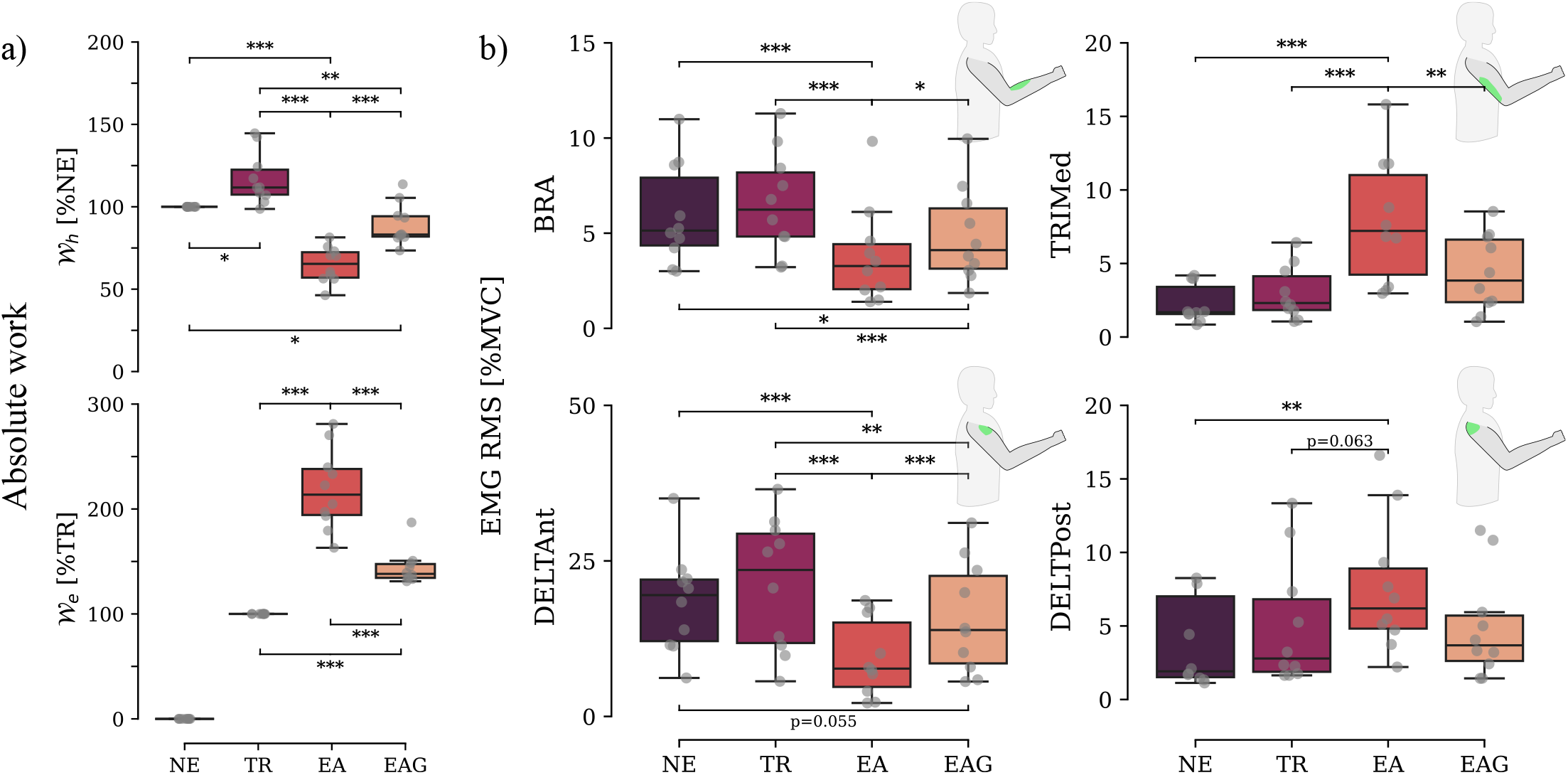
Human energetic expenditure in the different conditions. **a)** Normalized mechanical work of the human 𝒲_*h*_ and exoskeleton 𝒲_*e*_. **b)** Normalized RMS of the four main EMG signals used to estimate human joint torques. BRA and DELTAnt are the main contributors to shoulder and elbow flexion in the task, and TRIMed and DELTPost are representative of elbow and shoulder extension.

This analysis showed that both myocontrollers (EA and EAG) increased the contribution of the exoskeleton when compared to transparency (TR). This trend was observed for all phases of the proposed task, with a statistically significant main effect of the controller on the total absolute mechanical work (*p <* 0.001, *F* = 28.4, *η*^2^ = 0.76). More precisely, the EA control scheme allowed for a significant reduction in human work compared to the NE and TR conditions with a large effect size (*p <* 0.001, Cohen’s *d >* 1) and a medium effect size for EAG (*p <* 0.001, *d* = 0.67). The EAG controller also decreased energy consumption with a moderate effect against NE (*p <* 0.05, *d* = 0.36) and a large effect size against TR (*p <* 0.001, *d* = 0.88). In sum, the human energetic expenditure was overall lower with the EA and EAG controllers, being the lowest in the EA condition.

While the absolute mechanical work is a clear indicator of energy expenditure, it does not draw the complete picture of muscle effort allocation during the task. This allocation is described in Fig. 3b, with a full decomposition per muscles and movement phases available in supplementary materials. It highlights an overall increase in the activity of elbow and shoulder muscles when interacting with the exoskeleton during transparent control, which is consistent with previous studies [7]. In line with the results on mechanical work, this analysis shows that the EA and EAG controllers decreased the activity of the elbow and shoulder antigravity muscles compared to the TR controllers. This result was expected given that antigravity muscles are the main actuators during lifting tasks. Conversely, this assessment shows that EA increased the activity of the shoulder and elbow extensors. However, this increase remained limited from an energetic standpoint, as these muscles were not heavily involved in the task.

Further statistical analyses revealed significant variations in muscle activation patterns (*p <* 0.05, *F >* 5.66, *η*^2^ *>* 0.38) between the different controllers, expect for the posterior deltoid. Focusing on the flexor muscles, EA significantly reduced activation of the brachialis, brachioradialis and anterior deltoid throughout the movement cycle compared to NE and TR, with a large effect size (*p <* 0.001, *d >* 0.85). EAG resulted in an significant decrease in the activation of the brachialis and anterior deltoid compared to TR (*p <* 0.01, *d >* 0.82), althoug less effective for the brachioradialis (*p* = 0.008, *d* = 0.69). Overall, compared to TR, both EA and EAG reduced flexor muscles activation. Regarding extensor muscles, the EA controller caused a significant increase in activity for all three triceps heads compared to the other conditions (*p <* 0.05, *d >* 0.8). Although this was not significant for the posterior deltoid (*p* = 0.077), an increase in its activation was also induced with a large effect size (*d >* 1). Conversely, EAG did not increase extensor muscles’ activation compared to NE and TR. In conclusion, both the EA and EAG controllers reduced flexor muscle activation compared to TR and NE, with EA being the most effective. However, unlike EA, EAG successfully avoided to increase extensor muscle activation while achieving this reduction.

Results from muscle activation and mechanical work demonstrated that less effort was required when assistance was provided. However, as previously stated, effort should not be the only performance metric used to validate the relevance of a physical assistance. Although often overlooked, assessing the effect of the controllers on other components of human motor control is critical. For example, if an assistive control is too slow, it can result in unnecessary energy expenditure due to a misalignment with the user’s internalized cost of time [42], thus hindering acceptability. Below, we provide an in-depth assessment of the impact of different controllers on human motor control through kinematics, and analyze the self-reported ergonomics of the interaction.

### 2.2 Kinematics are more natural with uncompensated gravity

We first qualitatively assessed the effect the effects of the different conditions on human kinematics through shoulder and elbow joint trajectories. The average participants’ trajectories are reported in Fig. 4.

**Figure 4:**
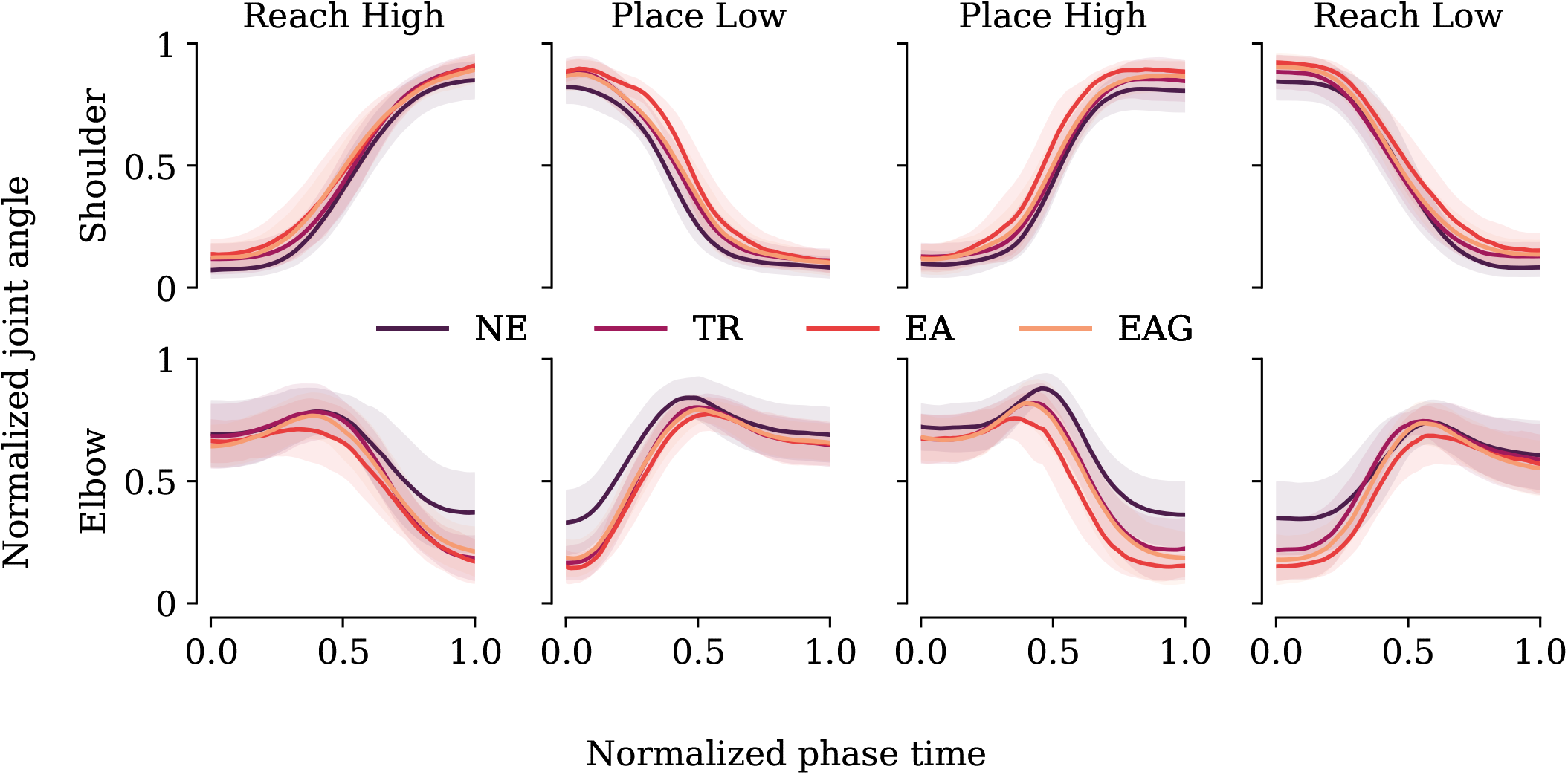
Impact of the exoskeleton on joint trajectories. Normalized joint angle during each phase of the movement. Each colored line correspond to one condition.

Compared to NE, the exoskeleton clearly influenced the user’s movements for both the shoulder and elbow joints during each phase. Setting aside any possible postural changes–necessarily limited due to the connection with the robot–comparisons between TR, EA and EAG suggest that the EA condition tends to systematically impact baseline human trajectories in the exoskeleton. In addition to being closer to TR, trajectories performed while interacting with the EAG controller were also closer to the TR baseline than those performed with EA, suggesting an overall smaller impact of this controller on human kinematics.

To validate these qualitative observations, we then computed several usual kinematic parameters that could indicate differences in the integration of the exoskeleton behavior by the human. Specifically, we analyze the movement duration, distance traveled by the hand and movement smoothness in Fig. 5, with a detailed version for each phase of the task provided in the supplementary materials.

**Figure 5:**
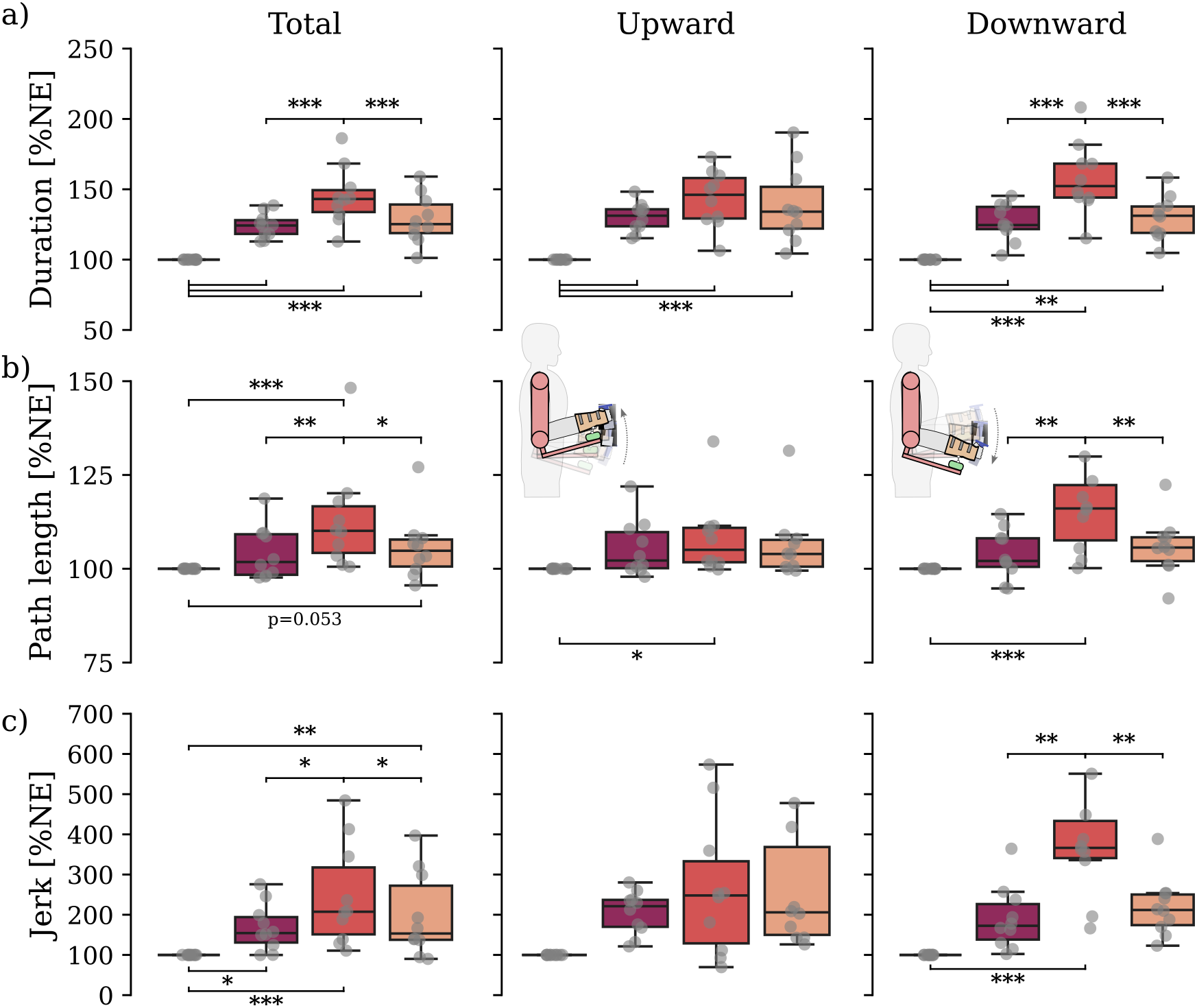
Impact of the exoskeleton on motor control parameters. For all the metrics, phases are regrouped by direction (*i*.*e*. Reach high and Place high are upward). **a)** Normalized duration (in % of the value obtained for the NE condition). **b)** Normalized total path length (in % of the value obtained for the NE condition). **c)** Jerk smoothness of movement profile. ***: p < 0.001, **: p < 0.01, *: p < 0.05

The findings presented in Fig.5a indicate that movements executed within the exoskeleton were typically slower compared to those executed outside of it (*p <* 0.001, *F* = 25.8, *η*^2^ = 0.74). Specifically, NE demonstrated a globally significantly shorter duration (*p <* 0.001, *d >* 1) relative to all other conditions. While no significant differences were found when considering upward movements, the EA controller resulted in the longest duration when considering downward movements, showing a significant increase compared to EAG and TR with a very large effect size (*p <* 0.001, *d >* 1). Interestingly, no significant difference was observed between EAG and TR for either direction (*p >* 0.47, *d <* 0.26). Overall, the exoskeleton slowed down human movements, particularly with the EA controller, which resulted in the slowest downward movements. The EAG controller did not lead to slower movements compared to TR.

The results depicted in Fig.5b show a significant main effect of the controller on the path length (*p <* 0.001, *F* = 9.9, *η*^2^ = 0.52). Specifically, while there were no differences in path length for upward (i.e. *reach high* and *place high*) movements, EA showcased significantly longer paths than the other three conditions for downward (i.e. *reach low* and *place low*) movements (*p <* 0.006, *d >* 0.56). As for movement duration, no differences were found between EAG and TR.

Similar differences were observed in terms of movement smoothness, as illustrated in Fig. 5c, with a main effect of the controller on the dimensionless jerk (*p <* 0.001, *F* = 11.7, *η*^2^ = 0.565) (see [46] for the definition). When considering downward movements, the EA conditions resulted in significantly lower smoothness compared to all other conditions (*p <* 0.006, *d >* 1), suggesting that users had difficulties to plan their actions as unconstrained human movements are known to be smooth [47, 48]. Similarly to the other parameters, we found no significant differences between EAG and TR, and upward movements did not result in significant differences among conditions.

These results show that both EMG-based assistance methods reduced effort, with contrasted impacts on human motor control. Specifically, the traditional EA method aiming to fully compensate human inputs, led to degraded kinematic performance, with slower, longer, and less smooth movements. However, these findings are not sufficient to validate our effort-sharing approach. When designing physical assistance controllers, user feedback regarding the perceived cognitive and physical ergonomics is critical. Here, we completed our analysis by asking participants to mark the different controllers on ergonomic scales, as reported in what follows.

### 2.3 Subjective perception favors uncompensated gravity

After each set of 25 cycles in a condition, participants were required to complete a questionnaire related to task perception that was adapted from NASA-TLX [49, 50] (see section 4.5.1). The results are summarized in Fig. 6.

**Figure 6:**
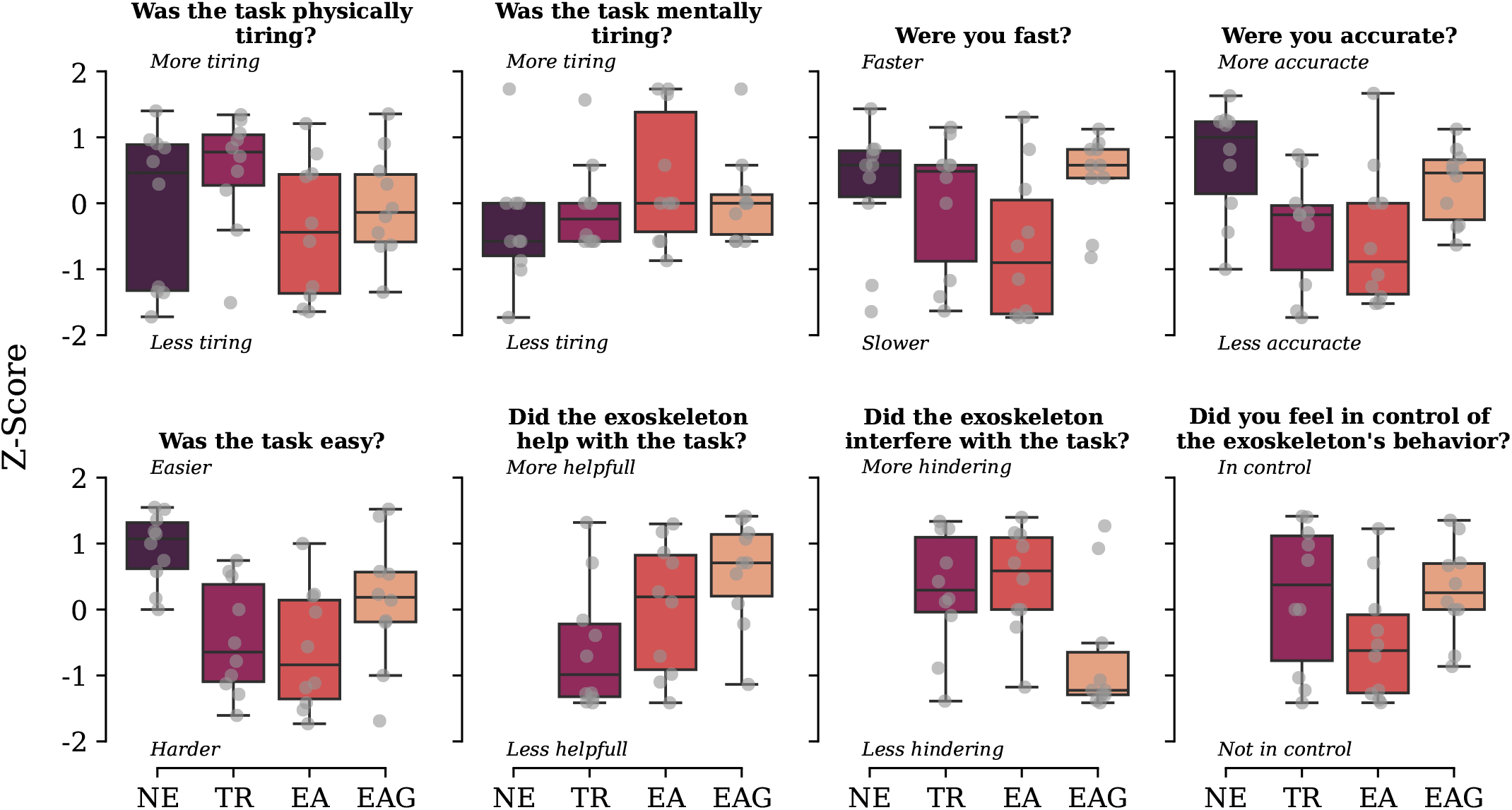
Z-scores of the questionnaire marks reported under different controllers.

Although there was considerable variability among participants, distinct patterns emerged depending on the controller. Participants found the task to be less tiring when using the exoskeleton with assistance control, with the EA controller being slightly less tiring on median when compared to EAG. However, participants reported feeling more mentally exhausted after using the EA controller. This was supported by other questions, indicating that while EA felt slightly less physically tiring, it also felt slower and less accurate. Consistently, participants felt that EA and TR hindered the completion of the task, and EA did not provide the same level of control over the exoskeleton when compared to both TR and EAG. EAG showed improvements over EA and TR for all those metrics, with a perceived velocity comparable to NE and an improved perceived accuracy when compared to TR and EA. While the EAG controller was still perceived as more challenging than NE, it remained a clear improvement in terms of both cognitive and physical ergonomics.

## 3 Discussion

In this study, we introduced two myocontrollers aiming to provide a versatile physical assistance seamlessly integrated by users. The first controller (EA) aimed at minimizing the user’s muscle activity using a proportional-integral scheme, which resulted from the combination of two existing approaches [9, 32]. Building on this foundation, the second controller (EAG) aimed at implementing a more intuitive physical assistance by leaving the arm’s gravity-related efforts being dealt by to the user. This was based on the fact that humans exploit gravity when controlling their vertical movements [37, 38], a current motor control knowledge which was also observed when interacting with an active exoskeleton [10]. The controllers were validated through comparisons against a baseline transparent control (TR) condition while performing a multi-joint displacement task, carrying an unknown load and inspired by movements observed in logistics.

As expected, both myocontrollers allowed to significantly decrease the energetic cost of the task when compared to transparent control. This reduction was particularly pronounced for flexor muscles such as the brachialis and anterior deltoid. However, when fully compensating for user effort, it was associated with unusual and counterintuitive muscle activation patterns of the extensor muscles during downward movements, leading to significantly decreased movement intuitiveness. This phenomenon was far less pronounced with uncompensated arm’s gravity, enabling more intuitive downward movements with kinematics closer to the baseline, albeit with a smaller reduction in energy expenditure. Importantly, when the user’s effort is fully compensated, the aforementioned impacts on kinematics may lead to decreased acceptability and unwanted interaction forces, as it is known that humans are reluctant to modify their movement velocity, in particular towards slower movements [42, 51]. Two main reasons may explain why uncompensated gravity torques are beneficial. First, the exoskeleton may be perceived as a disturbance; therefore, providing less assistance may result in less constrained movement, giving the user more freedom to impose their own kinematics. Second, not compensating for a portion of gravity-related efforts allows the users to use them to accelerate when moving downwards and decelerate when moving upwards, which more closely corresponds to usual human motor patterns [38,40,41] and may limit the required motor adaptations. Overall, we exhibit a trade-off between the human ability to seamlessly interact with EMG-based assistive exoskeletons and the level of provided assistance, higher assistance leading to lower interaction quality when impeding usual motor control patterns.

One of the key features of myocontrollers is that they do not need an *a priori* knowledge about the task, whether it involves manipulating objects with an unknown mass or following a desired path, which is usual in industry (*e*.*g*. in logistics warehouses [43]). These controllers dynamically adjust based solely on real-time EMG signals, allowing to obtain a more versatile assistance, where the system accounts for human movement variability and changing dynamics by design, without requiring a task-specific calibration. Conversely, traditional control systems rely on predefined models or demonstrations of the user’s intended path [11,16,17,52] or the characteristics of the object being manipulated [8, 30].

While the present study exhibits critical results regarding human motor control strategies for future EMG-driven exoskeleton assistance, some limitations remain. In particular, although we validated our method in a multi-joints setting, other degrees-of-freedom actuated by deep muscles such as shoulder internal/external rotations are very difficult to assist using surface EMG. More fundamentally, human muscle recruitment strategies are known to be variable over time [53], and EMG signals are highly impacted by muscle fatigue [54], prolonged wear [20] and electrode shifting [55], which phenomenons have to be considered to ensure the robustness of the assistance to the passing of time. Finally, there is a lack of common benchmarking tasks and exoskeletons to evaluate physical assistance controllers, which is even more problematic for EMG-based assistance where the sensors quality can largely impact the robot’s behavior. An example of this are the two studies we used to build our controllers: Lotti et al. [9] evaluated their method on horizontal pick-and-place tasks with an elbow assistance, while Treussart et al. [36] proposed a static gravity compensation with a different EMG system. It is unclear how such experiments could be compared together or to our more advanced task, warranting a standardization for the evaluation of EMG-based physical assistance, and more generally of upperlimb exoskeleton controllers. Our findings also highlight the need for further research on the proposed controllers. These were tested without any modification to the assistance or gravity gains (*i*.*e. α*_*a*_ = 1 and *α*_*g*_ = 1). Future research should focus on continuously adapting these gains with respect to kinematics to ensure both intuitiveness and effort reduction. Finally, control schemes must be tested in practical environments, such as industrial settings or neurorehabilitation programs to verify their relevance to a broader user base and population with specific activation patterns. This approach would address a significant shortcoming of laboratory experiments, in which participants have limited exposure to the exoskeleton [56]. Extended use in real-world scenarios could enhance user adaptability and influence energy expenditure and other kinematic criteria.

This work paves the way for more intuitive human-machine interaction in the future, particularly with regard to exoskeletons in assistive setups. These results clearly demonstrate that integrating and fusing neuroscientific insights into robotics controllers will enhance the effectiveness of assistive and medical devices. Despite remaining challenges, the myocontroller we introduced is a promising venue for exoskeleton assistance, leveraging the direct measure of user intent allowed by EMG sensors and providing adaptive and intuitive support without requiring prior knowledge of the task at hand.

## 4 Material and methods

### 4.1 Material and Participants

#### 4.1.1 Participants

10 healthy right-handed subjects (7 males, age 27.8 *±* 6.8 years, height 175 *±* 8 cm, weight 72.4 *±* 10 kg) took part in the experiment. The experimental protocol was approved by the ethics committee of Université Paris-Saclay (CER-PS-2021-048/A1). Participants signed a written informed consent form before starting the experiment.

#### 4.1.2 Exoskeleton

The ABLE4 exoskeleton [57] was used in the experiments. This highly backdrivable exoskeleton includes four actuated degrees of freedom (DoF), one for elbow flexion/extension and three for shoulder rotations. In these experiments, the elbow and shoulder flexion/extension were used, while the other two DoF were mechanically locked to constrain motion in the sagittal plane as in previous works [7, 45]. The exoskeleton was connected to the user through a self-aligning physical interface, maximizing comfort, designed and evaluated in prior studies [44, 45]. This interface provides a large contact surface and includes a passive spherical joint, torques from overconstraints. The interaction forces were recorded using a six-DoF force-torque (FT) sensor (1010 Digital FT, ATI, Apex, USA). Angular positions, speeds and interaction force were synchronously recorded at 1 kHz. The control schemes of the exoskeleton varied depending on the experiment (calibration or assistance) and condition (transparency or myocontroller), as detailed in Section 4.2.

#### 4.1.3 Motion capture

The kinematics of both the participant and the exoskeleton were captured at 100 Hz using 10 Oqus 500+ infrared cameras (Qualisys, Göteborg, Sweden). Reflective markers were attached to specific landmarks of the exoskeleton, allowing to extract segment positions and orientations. For the participant, the positioning of the reflective markers varied depending on the phase of the experiment (see Fig. 7.a and supplementary materials). First, markers were placed on the acromion, the seventh cervical vertebra, the sternal ends of the clavicles, and the styloid processes of the ulna and radius, as well as the lateral and medial epicondyles to extract each participant’s anthropometrics (see 4.2.2). Then, only markers on the acromion, clavicles, and vertebrae remained, while the others were removed as they were hidden by, or interfering with, the exoskeleton. Full upper-limb kinematics were determined using these remaining markers and those on the exoskeleton. An illustration of reflective makers placement is available as supplementary materials.

**Figure 7:**
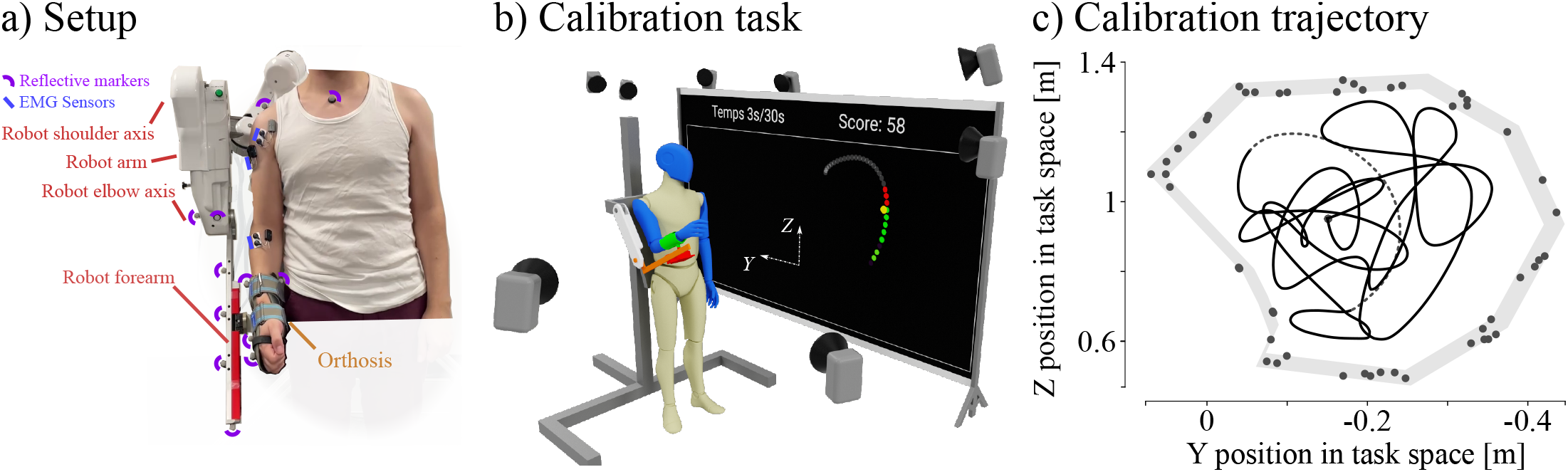
Detailed experimental setup and calibration task. **a)** EMG sensors and reflective markers were placed on relevant anatomical landmarks. **b)** For the calibration, the exoskeleton was laterally placed to a projected screen. Participant had to follow the trajectory displayed on the screen. **c)** Calibration trajectories were generated using b-splines. A cloud of spline nodes was generated around the task space, allowing the generated trajectory to maximize its exploration.

#### 4.1.4 Electromyography

Electromyography (EMG) signals were captured at 2 kHz using wireless MiniWave sensors (Cometa, Bareggio, Italy). Eight muscles that have been shown to provide an optimal tradeoff between the number of sensors and accuracy for sagittal plane movements [29] were recorded (see Fig. 7.a and supplementary materials): the anterior, posterior, and medial deltoids (DELTAnt, DELTPost, DELTMed), the brachioradialis (BRD), the brachialis (BRA), and the long, medial, and lateral triceps (TRILong, TRILat, TRIMed). Sensors were positioned as recommended by previous studies [58, 59]. An illustration of EMG sensor placements is available in supplementary materials. Signals were first band-pass filtered (Butterworth, fourth-order, cutoff frequencies {20, 450} Hz), before being centered and rectified. Their envelope was then extracted using a low-pass filter (Butterworth, fourth-order, cutoff frequency 3 Hz), as recommended in previous studies [29, 60]. Lastly, the signals were normalized using maximum voluntary contraction (MVC) values measured during dedicated trials.

### 4.2 Experimental procedures

The experimental procedure included several phases conducted sequentially after equipping the participants with the reflective markers and EMG sensors. It was organized as follows: (i) measurement of the anthropometric data and maximum voluntary contraction, (ii) measurement of training EMG and torque data, followed by a 10 min break for the EMG-to-torque model parameters optimization, and (iii) execution of the experimental task with one baseline condition without exoskeleton (NE) and three tested controllers (TR, EA and EAG).

#### 4.2.1 Anthropometrics and MVC

##### Anthropometrics

Initial anthropometric measurements were conducted using the reflective markers positioned on anatomical landmarks as outlined in the materials section (see 4.1.3). Participants were requested to enter the motion capture area, where a static pose was recorded. Subsequently, an existing upper-limb OpenSim model [61, 62] was adjusted to align the joints with the positions of the reflective markers, allowing to adapt limb dimensions and masses.

##### Maximum voluntary contraction (MVC) trials

MVC trials were performed in four positions, each maximizing the contraction of a muscle group: elbow flexion, elbow extension, shoulder flexion, shoulder extension. The contraction lasted three seconds and was performed twice with a resting period in-between. The maximum values recorded across all trials were then used to normalize EMG signals.

#### 4.2.2 Calibration experiment

This experiment aims at identifying the parameters of the EMG-to-torque model. During this phase, a preexisting calibration framework [28], that has provided clean dataset for EMG-to-torque models calibration [63], was used. Participants were asked to perform a dynamic tracking task, consisting in following a randomly generated trajectory while the exoskeleton provided a viscous resistance (see Eq. 5 and Fig. 7.b). Five 30*s* trajectories were used to gather calibration data. Each trajectory was randomly generated to provide a wide range of kinematic and dynamic situations, covering the whole workspace to ensure a robust calibration [28](see Fig. 7.c). To do so, the target trajectories were parameterized by B-splines, for which the control points were randomly placed in task space. For each calibration trial, a hundred candidate trajectories were generated, and the one maximizing the mechanical work and the workspace coverage was selected. The trajectories were displayed on a screen positioned laterally to the exoskeleton, with visual feedback of the participant’s hand position.

Arm kinematics, human-exoskeleton interaction efforts, and EMG signals of upper-limb muscles were recorded. Shoulder and elbow joint trajectories and torques were extracted via inverse kinematics and dynamics by incorporating the recorded positions of the reflective markers and the efforts in OpenSim. The resulting dynamic data were then used to calibrate the EMG-to-torque model as described in Section 4.3.

#### 4.2.3 Assistance task

To assess the effects of the tested control methods (TR, EA and EAG) on human effort, behavior and perception, a pick-and-place task was designed. This task mimics what would happen in an ecological situation, where movement phases can be performed with and without external loads.

The load was a 1.5 kg steel weight attached to a custom 3D-printed handle. The handle was specifically designed to facilitate the carrying of the weight with limited grasp force (i.e. the top of the handle rested on the thumb phalanges and the distal end of the second metacarpal bone), thereby limiting cross-talk from the forearm muscles to the brachioradialis EMG sensor. Two platforms with adjustable positions and heights served as stations for picking and placing the weight (see Fig. 2.b). The first platform was positioned so that the weight could be picked up with the upper arm in a vertical position alongside the body, and with the elbow bent at a 90^°^ angle. The positioning of the second platform was determined through an iterative process. Initially, it was placed at shoulder height and at a forearm’s length from the first platform. Minor adjustments in platform position were then applied to make sure participants were able to reach every position of the tasks without being limited by the reachable area provided by the exoskeleton

The task, which is illustrated with a supplementary video, involved a sequence of shoulder and elbow movements in a parasagittal plane divided into four phases. At the beginning, the weight was placed on the upper platform, while the participant’s hand was at the starting position, which corresponded to the picking point on the lower platform. First, the participant had to reach for the weight, referred to as reach-high (RH) movement phase. Second, the participant picked the weight and placed it at the lower position, referred to as place-low (PL) movement phase. Third, the participant picked the weight again and placed it back at the upper position, referred to as the place-high (PH) movement phase. Finally, the participant moved their hand back to the starting position, referred to as the reach-low (RL) movement phase. This sequence of four operations enabled us to compare the effects of assistance between upward and downward movements, both with and without weight. After completing the four phases, both the participant’s hand and the weight were back to their original positions, allowing the participant to start a new movement cycle. For each of the four conditions, this cycle was performed 25 times, with a four-minute break between conditions, inducing a total of 100 cycles.

The participants performed the pick-and-place task under four conditions as follows: (i) without exoskeleton (NE), with a light splint reproducing its orthosis, serving as a familiarization period, (ii) with transparent control (TR), to establish a baseline within the exoskeleton, and (iii) with the myocontrollers (EA and EAG), performed in a random order subject-wise.

### 4.3 Mapping muscles activities to joint torques

Providing an accurate and timely estimation of the instantaneous joint torques of the user is a foundational component of a continuous assistive control framework. This basic component requires (i) a model mapping processed EMG signals to joint torques, and (ii) an individualized calibration of this model’s parameters, both described below.

#### 4.3.1 EMG-to-torque model

A non-linear mapping model (NLMap) [28] was used to transform EMG signals into joint torques. It is composed of a non-linear activation computed from EMG then linearly mapped to torques, which was shown to outperform more complex models describing explicitly the neuromusculoskeletal (NMS) system [28], with a shorter calibration time. The non-linear activation function, inspired by the shape factor of NMS models [64], maps the EMG signals **m**(*t*) ∈ ℝ^8^ to muscle activations:

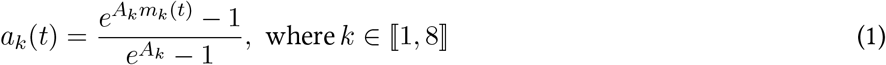

with *a*_*k*_(*t*) the non-linear activation of the *k*^*th*^ muscle and *A*_*k*_ the corresponding shape factor. These activations are then linearly mapped to joint torques:

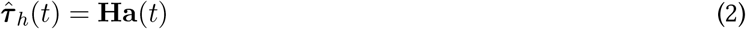

where 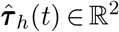 is the vector of estimated human joint torques, **H** ∈ ℝ^2*×*8^ the linear mapping matrix, and **a**(*t*) ∈ ℝ^8^ the vector of muscle activations.

#### 4.3.2 Model calibration

##### Parameters optimization

Importantly, the model of Eq. 2 needs to be calibrated for each user. In the present study, calibration data were collected during a dedicated calibration experiment (see 4.2.2), allowing to obtain human torques **T**_*c*_ ∈ ℝ^2*×N*^, from inverse dynamics, and associated EMG signals **M**_*c*_ ∈ ℝ^8*×N*^, with *N* the number of calibration samples. The NLMap model requires a bi-level optimization to identify (i) the shape factors *A*_*k*_, and (ii) the weights of the linear mapping matrix **H**, which was achieved using a single objective genetic algorithm [25, 65]. This algorithm involves an iterative process aiming at finding a set of parameters, referred to as a chromosome, that minimizes the squared torque estimation error:

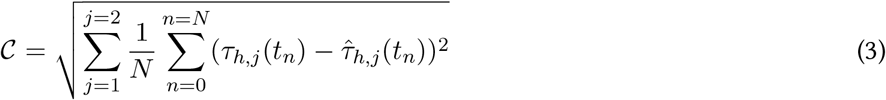

where ***τ***_*h*_ is the measured human joint torque extracted from **T**_*c*_. During each iteration, a chromosome containing a set of shape factors was generated through mutation and cross-over. Subsequently, the non-linear activations **a** were computed and compiled in a matrix **A**_*c*_ ∈ ℝ^8*×N*^, which was used to identify **H** by solving:

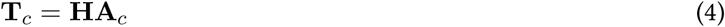

which was done using Matlab’s *mvregress* function (MathWorks, Natick, MA, USA). The algorithm terminates either when the maximum number of iterations is reached or when the cost function stalls, allowing to obtain the optimal shape factors and linear mapping coefficients with respect to Eq. 3.

##### Exoskeleton control

During the calibration experiment, the objective was to maximize the variations of activity of the measured muscles while moving in the whole workspace, for both upwards (*i*.*e*. against gravity) and downwards (*i*.*e*. in the same directions as gravity) movements. Here, accounting for direction is important as it is known that, during downwards movements, humans start by deactivating antigravity muscles (used to move upwards) instead of activating gravity muscles (*i*.*e*. used to move downwards) [10, 41]. Therefore, to make sure that the calibration data contains extensor activations, an open-loop gravity compensation of the exoskeleton and an end-effector viscous resistance were applied:

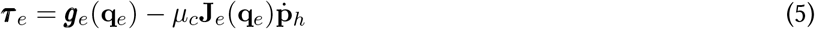

where ***τ***_*e*_ ∈ ℝ^2^ are the exoskeleton control torques, **q**_*e*_ ∈ ℝ^2^ its angular positions, **J**_*e*_ its Jacobian matrix, **g**_*e*_ its gravity model, and 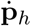 the participant’s hand velocity. The viscous load was personalized for each subject, ensuring a moderate effort on the Borg’s scale.

### 4.4 Exoskeleton controllers

In the present paper, three controllers were tested and their effects on the human performance and behavior in the main task were assessed (see Section 4.2). The controllers may be defined as follows: (i) a transparent controller (TR), minimizing the interaction efforts, to serve as a baseline, (ii) an assistive myocontroller minimizing the estimated human torques (EA), and (iii) a second assistive myocontroller which did not compensate for the gravity component of the user’s arm (EAG). These controllers were implemented as described below, and a global block diagram is pictured on Fig. 2.c.

#### 4.4.1 Transparent controller (TR)

The transparent controller aims at following human movements while minimizing human-exoskeleton interaction efforts. Similar to previous works on this exoskeleton [6,7], a combination of open-loop compensation of the exoskeleton’s weight and a closed-loop minimization of interaction efforts using a proportional-integral correction were used. This correction was used on the interaction forces measured via the FT sensor, **F**_*i*_. Note that the torques measured by the FT sensor were not used because of the passive ball joint in the human-exoskeleton interface [44, 45]. The exoskeleton control torque ***τ***_*e*_ was therefore:

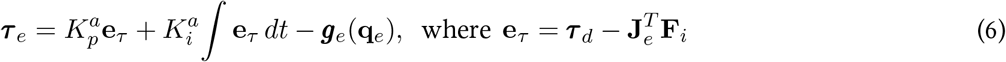

with ***τ***_*d*_ = **0** the null desired torque for admittance control, 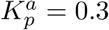 and 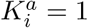 the proportional and integral gains, and ***g***_*e*_(**q**_*e*_) the open-loop compensation of the exoskeleton’s weight based on joint positions **q**_*e*_.

#### 4.4.2 Myocontrollers

The assistance myocontrollers were built on top of the TR controller, with ***τ***_*d*_ determination based on human joint torques (see Eqs. 1,2). Compared to other kinematic-based [52] or task-based controllers [8, 11], EMG-based controllers present a functionnal advantage in terms of the necessary *a priori* knowledge. In particular, for tasks involving a direct interaction between the user (*e*.*g*. its hand) and its environment (*e*.*g*. an object), no *a priori* knowledge about the trajectory or object mass is necessary, as explained below.

##### Absence of trajectory and mass knowledge

First let’s consider the full dynamics of the human’s upper limb in the exoskeleton:

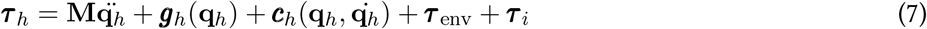

where ***τ***_*h*_, ***τ***_env_ and ***τ***_*i*_ are the torques respectively corresponding to human muscles, to the interaction between the human and its environment (*e*.*g*. an object), and to the interaction between the human and the exoskeleton (*i*.*e*. the assistance torques). **M, *g***_*h*_, and ***c***_*h*_ respectively correspond to the human mass matrix, the gravity and the centrifugal and Coriolis torques vectors, with **q**_*h*_ the joints coordinates. In the case of a classical dynamic compensation, an open-loop control would aim at setting the exoskeleton torques so that:

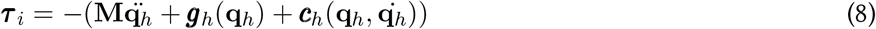

letting the remaining human effort to be ***τ***_*h*_ = ***τ***_env_. With the addition of trajectory prediction based on task learning [52] and a prior knowledge of the interaction with the environment, one can even attain a net null human effort by estimating ***τ***_env_. This method cannot however achieve general movement assistance as a prior learning remains necessary for any new or sufficiently different task.

The proposed myocontrollers on the other hand operate in a closed-loop, directly using the estimated human torques ***τ***_*h*_ as the feedback. As a consequence, the correctors will directly ensure that ***τ***_*h*_ converges to zero rather than relying on the dynamic model of the human effort. Therefore, during assistance, and considering a sufficiently accurate human torque prediction, this convergence (*i*.*e*. ***τ***_*h*_ → 0), will result in the exoskeleton torque converging to the equivalent dynamics and interaction:

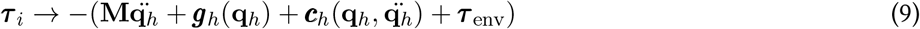

It is important to note that this convergence is blind to the terms of the equation, and thus model-free. It is therefore not possible to extract the value of a single term without prior knowledge about the others. Still, the resulting control successfully compensates the full dynamics and the interactions of the upper limb without information about the trajectory or the environment. The implementation of the two versions of this control approach is described below.

##### Basic EMG-assist controller (EA)

Starting from the EMG-to-torque model, and once calibrated, the estimated torques were (i) filtered using a dead zone, with threshold T = 1 N.m in order to manage minor fluctuations during low torques situations [32], and (ii) multiplied by an assistance gain *α*_*a*_ setting the proportion of task torques compensated by the exoskeleton, resulting in the following human torque 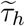 to compensate:

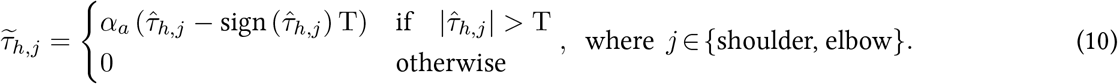

Following the phasic and tonic equivalence of human movement, a proportional-integral corrector scheme was used to compute the desired interaction torques:

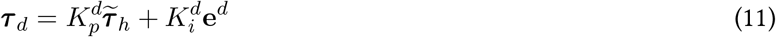

with 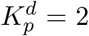 and 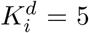 the proportional and integral gains of the assistance torque corrector. Importantly, the role of the integrator in the present controller is quite different when compared to its usual role. Here, it acts as a memory of efforts, allowing to solve the paradox of purely proportional assistance methods [9, 33–35], where more assistance entails less EMG signal, leading in turn to less assistance. Conversely, by considering the human inputs as a signed error **e**^*d*^ accumulating through time, it is possible to memorize the current requested effort and provide a constant assistance, even when EMG activity decreases. For instance, in a task requiring to hold a mass at a given posture, the assistance will initially rely on its proportional component, but rapidly, as the **e**^*d*^ term increases, the proportional term will become less important because the EMG activity will decrease, and the assistance will rely primarily on its integral component. The values of 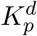 and 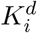 were chosen so as to maximize the rate of transfer from the proportional to the integral components of the assistance, thereby allowing to minimize the need for EMG activity to perform the assistance. This effort memory is computed as follows:

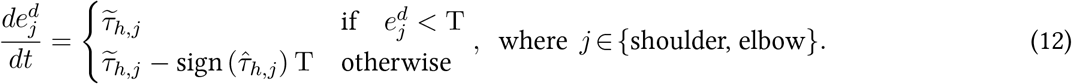

The resulting desired interactio n torque ***τ***_*d*_ is then used in the the previously described admittance scheme.

##### EMG-assist controller with intuitive gravity integration (EAG)

As an alternative to the previous control strategy that only considered muscle activities, a controller that accounted separately for gravity-related torques was developed. The hypothesis was that, since these torques are an ubiquitous component of our environment that have been shown to be very well optimized and integrated by the central nervous system in motion planning [37], not accounting for them in the myocontroller may result in less intuitive interactions with the exoskeleton. Specifically, after memorizing the level of effort required to maintain a posture with the integral term, the user will not be able to simply reduce the activity of antigravity muscles to initiate downward movements as expected [10, 41]. To address this issue, the negative arm gravity torques were added to the estimated human effort from EMG signals, thereby removing them from the desired assistance torque:

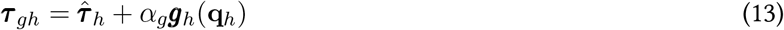

with *α*_*g*_ the gravity gain setting the proportion of the arm gravity torques uncompensated by the assistance. Specifically, *α*_*g*_ = 1 leads to a normal gravity context for the human’s arm, *α*_*g*_ = 1*/*6 leads to a gravity close to the Moon’s, and *α*_*g*_ = 0 leads to a complete compensation of gravity, as with EA. Here it is important to note that only the arm’s own gravity is uncompensated with the addition of this gravity term, any load carried through the hand remains assisted. The resulting human intention torque ***τ***_*gh*_ was then filtered and transformed into a control torque for the exoskeleton with the same deadzone and proportional-integral scheme as for the EA controller (see Fig. 2c).

### 4.5 Controller evaluation

#### 4.5.1 Ergonomic assessment

For each tested controller, participants were asked to provide a subjective assessment of the effects of the exoskeleton using a questionnaire adapted from NASA-TLX [49, 50] (see Table 1). For each questions, participants were asked to give an answer based on a continuous Likert scale (see table 2) that was displayed in front of them.

**Table 1.**
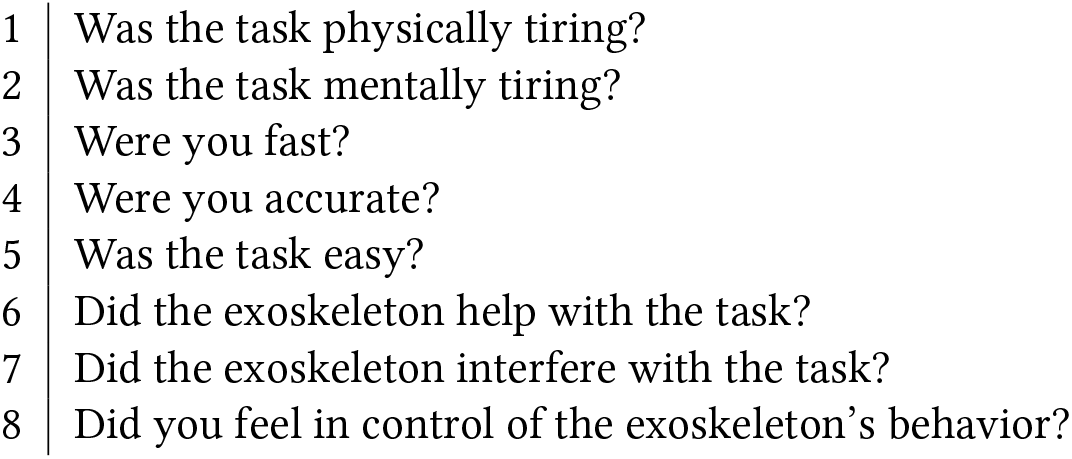
Subjective assessment questions. Translated from french to english. Those questions were asked after each condition. Questions 6 to 8 were not asked during the NE condition.

**Table 2.**
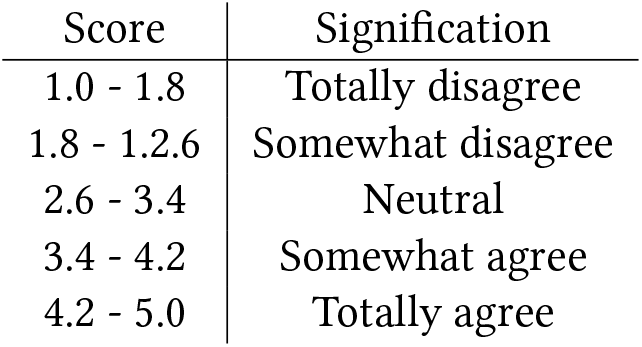
Likert scale. Participants were allowed to give a continuous value on the Likert scale, with a suggested signification of score ranges.

#### 4.5.2 Data processing

The data was segmented to isolate each phase of the movements performed by the participants in the main experiment (see Section. 4.2.3). Then, three criteria were used to assess the effects of the different controllers on human movement and effort.

##### Absolute mechanical work

The share of mechanical work performed by the exoskeleton and the human served as the primary criteria for evaluating the effect of the assistance provided by the exoskeleton. The absolute mechanical work expended by the human (*W*_*h*_) and the exoskeleton (*W*_*e*_) were computed as follows,

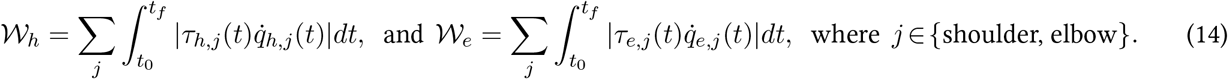

Similarly to all the subsequent metrics, the absolute mechanical works were computed for each segment related to a subject, a condition, a movement, and a phase of movement.

##### Muscle activities

The root mean square (RMS) values of each EMG signal were used as a secondary assessment of the human energetic expenditure in the task. They were computed as follows,

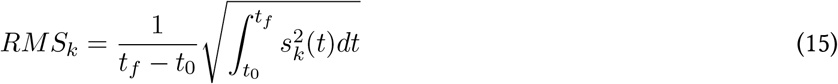

where *k* is the muscle identifier, *s*_*k*_(*t*) represents the filtered and normalized EMG signal, and *t*_0_ and *t*_*f*_ the beginning and end time of the considered movement phase..

##### Kinematics

Movement kinematics were assessed using both the cartesian position of the hand and the joint positions computed via OpenSim. Several criteria were used to assess behavioral changes associated with each tested controller compared to transparency and to the condition without exoskeleton. First, the duration of each phase of each trial was extracted, as well as its total duration. Then, the path length of the hand trajectory was computed for each trial/phase of trial as:

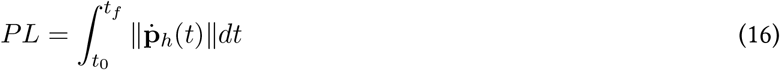

with **p**_*h*_ the cartesian position of the wrist, and *t*_0_ and *t*_*f*_ the start and end times of the given task segment. Finally, the dimensionless jerk, a measure of movement smoothness, was computed as:

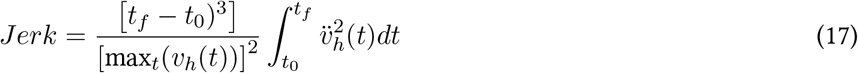

with *v*_*h*_ the norm of the cartesian speed such that 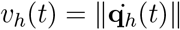.

#### 4.5.3 Statistical analysis

For each criterion, the median value for each participant was first computed before performing comparisons. The first ten movement cycles were excluded to account for adaptation [66]. To assess differences between conditions, a repeated measures analysis of variance (rm-ANOVA) was conducted with the condition as the within-subject variable and criterion value as the dependent variable. Mauchly’s test was used to check sphericity, and a Greenhouse-Geisser correction was applied if needed (*ϵ <* 0.75). Post-hoc analyses were performed after a Shapiro-Wilk test, to assess normality violations. Based on the results, either paired sample *t*-tests with a Holm-Bonferroni corrections, or Wilcoxon tests were conducted. For all tests, the significance level was set at *p* = 0.05 and Cohen’s *d* was used as a measure of the effect size.

## 5 CRediT authorship contribution statement

**Lucas Quesada**: Writing - Original Draft, Writing - Review and editing, Conceptualization, Methodology, Software, Formal analysis, Investigation, Data Curation, Visualization, Funding acquisition.

**Dorian Verdel**: Writing - Original Draft, Writing - Review and editing, Conceptualization, Methodology, Data Curation, Software.

**Olivier Bruneau**: Writing - Review and editing, Conceptualization, Supervision, Project administration, Funding acquisition.

**Bastien Berret**: Writing - Review and editing, Conceptualization, Supervision, Project administration, Funding acquisition.

**Michel-Ange Amorim**: Writing - Review and editing, Conceptualization, Supervision, Project administration, Funding acquisition.

**Nicolas Vignais**: Writing - Review and editing, Conceptualization, Supervision, Project administration, Funding acquisition.

